# Correlating amino acid profiles with improved expression capability of superior CHO based expression platforms

**DOI:** 10.1101/2024.08.16.608302

**Authors:** Vijeta Sharma, Richa Guleria, K. J. Mukherjee

**Affiliations:** School of Biotechnology, Jawaharlal Nehru University, New Delhi, India

## Abstract

A novel modified CHO cell line was constructed by expressing a three gene combination of Bclx(L), Aven and BECN1 which are critical targets of the apoptosis and autophagy pathways. This synergistic combination significantly improved cell viability as well as Rituximab (RTX) expression by 7.5 fold. A comparative amino acid analysis of the modified and control cultures both with and without RTX expression showed interesting changes in amino acid uptake rates. We first compared the utilization of amino acids in the control cells without RTX expression to the control cells with RTX expression where it declined significantly in the later half of the culture with the maximum decrease observed with tyrosine, which instead of uptake got secreted into the medium. The uptake of aspartic and glutamic acid also fell drastically during this period demonstrating the deleterious effect of RTX expression on cellular health. When the modified cells were used for RTX expression, there was conversely a slight increase in amino acid consumption in the later part of the culture with the maximum increase observed with alanine and tyrosine. The differential amino acid consumption rates provided us with an indirect measure of improved cellular health and ability of the modified cells to counter the cellular stress associated with RTX expression. Amino acid analysis could thus become a useful predictive tool to identify the critical features of better host expression platforms.

## Introduction

CHO cells are the backbone of the process for industrial production of therapeutically important proteins, like Rituximab (RTX), Bevacizumab and other blockbuster drugs like Tissue Plasminogen Activator, Erythropoietin and Factor VIII^1^. The upstream development of CHO cell lines to improve product quality and obtain high titers thus remains a major focus of research. A significant factor which critically impacts on production is the stress encountered by cells during the process of growth and product formation. This triggers complex responses which alter the cellular physiology and reduce product formation rates by affecting cell viability. Programmed cell death is one of the primary reasons behind the shortened product formation phase in a bioreactor^2–5^. Research has therefore been focused on blocking apoptosis pathways based on various knock-in^6–10^, knock-out^11–13^ and knock-down^14–16^ strategies. Since autophagy has also emerged as an important factor in reducing cell viability^17,18^, genes involved in this pathway have also been targeted for modification^19,20^. Pro survival role of autophagy during inevitable cell death is well established^21^, and therefore, synergistic strategies involving both apoptosis and autophagy genes have been used by researchers as an engineering strategy to increase cell viability^22,23^.

We decided to use a similar strategy to design an improved expression platform with simultaneous apoptosis and autophagy targeting in CHO cells. For this we chose Rituximab as a model protein, since it is a widely used antibody with applications in cancer therapy, auto immune disorders and transplant rejection. We took a previously untested, novel three gene combination of Aven, Bclx(L) and BECN1, which play critical roles in both apoptosis and autophagy and obtained enhanced cell survival as well as higher levels of protein expression.

While this result was important by itself, it allowed us to ask an equally relevant question viz. whether there exist associated phenotypic markers which correlate with this improved performance of the modified strain? Such markers could be identified by simply comparing the key metabolic features of the modified strain with the relatively poorly performing control cells. If this correlation could be validated it would provide a useful handle to predict and validate other better performing strains.

Such an exercise assumes significance given the fairly large number of potential combinations of gene knock-ins and knock-outs that can synergistically improve cellular health and productivity. A useful analytical tool for measuring cellular health is monitoring amino acid uptake rates which provide a glimpse of the molecular transformations taking place inside the cell^24–26^. We therefore decided to do comparative analysis of the modified and control strains with specific focus on amino acid consumption and/or secretion. Although studies have been done on the amino acid requirement of naïve and antibody producing CHO cells at various phases of growth^27,28^, amino acid analysis in correlation to cellular health has not yet been explored. To underpin the metabolic phenotypic changes multigenic cell engineering has been suggested by some researchers^29^. The goal of the present study was to use the comparative amino acid utilization profiles in order to identify key amino acid uptake rates which could serve as potential indicators of cellular health. Such a study could provide key guidelines in the identification as well as rational design of improved host expression platforms.

## Material and methods

### Cell line maintenance

CHO DG44 cells (*dhfr-*deficient) were provided by Dr. Lawrence Chasin, Columbia University, New York, USA. Cells were maintained in MEM α with 10% FBS. For each sub-culturing, 80-90% confluent cells were diluted in a new flask in pre-warmed complete medium to give a final cell density of 2 x 10^5^ viable cells ml^-1^. This was followed by incubation of the culture flasks in a CO2 incubator (Thermo Scientific, Massachusetts, USA) at 37°C with 80-90% humidity and 5% CO2.

### Construction of modified clones

The Bclx(L) and Aven genes were cloned into a bi-cistronic pBUD-EFGP plasmid (Addgene Plasmid #23027). The Bclx(L) gene was amplified from pMIG Bcl-x(L) (Addgene Plasmid #8790) using the primers Bclx(L)-F (5’-ATTTGCGGCCGCACCATGTCTCAGAGCAACCG-3’) and Bclx(L)-R (5’-GAAGATCTTCATT TCCGACTGAAGAGTGAGCCCAG-3’). The EGFP gene of the pBUD-EFGP plasmid was then replaced by Bclx(L) at the *BglII* and *NotI* restriction sites under the *EF1α* promoter. Similarly, using primers Aven_F (5’-GCGTCGACATCACCATGCAGGCGGAGCGA-3’) and Aven_R (5’-GCTCTAGAGGAAATCATGCTGTCCAACCAG-3’), the Aven gene was amplified from the SG5-Aven plasmid (provided by J Marie Hardwick, Johns Hopkins University, Maryland, United States) and cloned at the SalI-XbaI restriction sites of the pBUD-Bcl-x(L) plasmid under the CMV promoter (Figure S1(A) and S1(B)). The expression of both the genes was checked by western blotting after transient transfection into CHO cells (Figure S1C).

Since the plasmid having RTX (the protein of interest) had G418 as a selection marker, we amplified BECN1 gene (from pcDNA3-Beclin1 having G418 selection marker; Addgene: Plasmid # 21150) using primers BECN_F (5’-GCTCTAGAATGGAAGGGTCTAAGAC-3’) and BECN_R (5’-CGGGATCCTCATTTGTTATAAAATTGT-3’) and cloned it in pcDNA3-Hygromycin plasmid at the BamHI –XbaI site (Figure S2A and S2B). The expression was checked using Western blotting after transfection (Figure S2C). Stable cell line was generated by co-transfection of pBUD-Bclx(L)-Aven and pcDNA-Hygro-BECN1 using Lipofectamine 2000 using the manufacturer’s protocol. Stable clones were selected by limited dilution method.

### Construction of RTX expressing normal and engineered cells

The Bi-cistronic pcDNA 3.1(+)-G418 plasmid containing the DHFR gene plus the heavy and light chains of Rituximab (constructed in this study) was used for transient transfection of modified CHO DG44 cells as well as control cells. Cells were seeded in ultra low binding separate pre labeled T 25 flasks, 24 hours prior to transfection. On the day of transfection, DNA and bPEI in 2:3 ratio were mixed in 150 mM NaCl and incubated for 10 minutes. Cells were washed with 1X PBS and supplemented with CHO-S-SFM II media without antibiotic. The DNA-bPEI mixture was added to the cells and the flask was kept undisturbed for 24 hours in a CO2 incubator at 37⁰C with 5% CO2 and 90% humidity. The media was replaced by 7 ml MEMα with 5% fetal bovine serum and G418 antibiotic (Invitrogen, California, United States, cat. # ). The cells were counted every 24 hours and the culture media was frozen in liquid nitrogen for further analysis.

### Enzymatic assay(s)

The antibody concentrations in the culture supernatants were measured by ELISA using 96-well microtiter plates coated with a rat anti-rituximab antibody MB2A4 (Roche, Basel, Switzerland, cat. # MCA2260) and a Horseradish peroxidase (HRP)-conjugated detection antibody (Santa Cruz, California, United States, cat. # sc-2032).

### Quantitation of Apoptosis and Autophagy

Apoptosis was quantified using PI-Annexin V FITC based FACS assay. Briefly, cells were washed twice with PBS and twice with Annexin V binding buffer (Invitrogen, USA, cat. # 13246). The washed pallet was then resuspended in 100 µl of binding buffer and 2 µl of FITC tagged Annexin V (Invitrogen, USA, cat. # A13199) and 1 ul of 1 mg ml^-1^ PI (Sigma Aldrich, USA, cat. # P4864) was added to cells. After 10 minutes incubation in dark, cells were again washed with binding buffer and analyzed for fluorescence in flow cytometer. The qualitative estimation of apoptosis was also done by dual acridine orange/ethidium bromide (AO/EB) staining.

The autophagy assay was done by Western blotting using LC3 antibody (Sigma Aldrich, St. Louis, United States, cat. # L8918, Lot No. 013M4852V, ratio used 1:1000) and changes in the extent of autophagy were determined by Western blotting signal ratio between LC3-I and LC3-II. LC3-I (a cytosolic form of LC3) and LC3-II (a membrane associated phosphatidylethanolamine (PE) conjugated LC3-I) posses a molecular weight of 16 kDa and 14 kDa respectively and can be easily distinguished based on their differential mobility in SDS– PAGE. The level of LC3-II protein correlates with the number of autophagosomes in the cell. When cells undergo starvation for short periods, the amount of LC3-II increases with decrease in LC3-I levels, but if starvation persists for longer periods both LC3-I and LC3-II disappear in immunoblot assay^30^.

### Sample derivatization

Cell culture broth was collected for seven days at 24 hour intervals. For amino acid analysis, the lyophilized supernatant was derivatized (silylation) by adding N,N-dimethylformamide (Sigma Aldrich, USA, cat. # 227056) and MTBSFA (Tokyo Chemical Industry Co., Ltd., Tokyo, Japan, cat. # B1150) (1:1) followed by incubation at 85°C for 60 min. A standard solution containing all amino acids (at 2.5mM concentration for each amino acid) was also derivatized with MTBSTFA and was used for making standard calibration curve for each amino acid. D-norvaline standard solution (0.5 mM) was spiked in derivatized standards and biological samples (50 μL) enabling internal standardization for all amino acids. 1 µl of the derivatized sample was injected for GC MS analysis.

### Analytical methods

Glucose and lactate were analyzed using an HPLC system (1260 Infinity series, Agilent Technologies, USA) equipped with a refractive index detector following isocratic elution on a Aminex HPX-87H column (300 x 7.8 mm, 9 µm particle size; Bio-Rad Laboratories, CA, USA, cat. # 1250140) for a total run time of 20 min employing 5mM H2SO4 as mobile phase buffer. Flow rate was set at 0.5 mL min^-1^.

Amino acid analysis was performed using a GC-MS/MS instrument (Agilent Technologies, USA) equipped with an autosampler, a 7890B series gas chromatograph with a split/splitless injector and a mass selective detector coupled with an Agilent 7000B QqQ MS. The GC was operated in constant pressure mode (20 psi) with helium as the carrier gas and d5-glutamate as a standard for retention time locking. Agilent J&W DB-5MS column (30 m in length, 0.25 mm inner diameter, 0.25 μm film thickness with 10 m guard column) was used. The inlet pressure was 20 psi, temperature was set at 250°C, and injection (1 μL) was performed in the pulsed splitless mode. The GC oven was kept at 40°C for 2 min, and then raised to 320°C using a linear gradient of 30°C/min and finally held at 320°C for 4 min. The total run time was 30 min. The MS transfer line temperature was set to 280°C. Electron impact (EI) ionization was used with QqQ MS detection. Both HPLC and GC-MS analysis were performed in biological triplicates.

## Results

### Construction of RTX containing plasmid, transient transfection and expression

For expression studies we decided to use Rituximab as a reporter gene. This choice of the reporter was dictated by its therapeutic relevance and the fact that CHO cells are the most common hosts for production of monoclonal antibodies. The vector chosen for this was the bicistronic pcDNA3.1(+) Neo plasmid. The heavy and light chains of the RTX gene were synthesized separately and inserted under the CMV promoters between *NheI-EcoRI* and *NheI-XhoI*, respectively (Figure S3). Construction was confirmed by restriction digestion and sequencing. CHO cells were transiently transfected with this plasmid using an optimized protocol described earlier. The transfected cells were grown in stationary flasks in MEM alpha medium and RTX expression was confirmed by ELISA.

### Construction of modified CHO cells

As mentioned in material and methods section, the Bclx(L), Aven and BECN1 genes were PCR amplified from the respective plasmids. Restriction sites which were incorporated in the primers allowed us to insert these genes in the pBUD vector. The Aven gene was inserted downstream of the CMV promoter at the *SalI-XbaI* site and the Bclx(L) gene was under the *EF1α* promoter using *BglII* and *NotI* restriction sites (Figure S1(B)). The PCR amplified BECN1 gene was inserted into pcDNA 3.1 Hygro (-) plasmid at the *BamHI-XbaI* restriction site (Figure S2(B)). The control cells were co-transfected with these two constructs keeping the ratio 1:1 using Lipofectamine 2000 and stably transfected cells were selected by alternately exposing them to media containing zeocin (Invitrogen, USA, cat. # R25001) followed by hygromycin (Himedia Laborateries, Mumbai, India, cat. # A015). The stable cell pool from a single cell was created using limited dilution method and named DG44-Bclx(L)-Aven-BECN1. The expression of all the three genes from this single isolated pool was checked by Western blotting **(Figure 1)**. These modified cells were also transiently transfected with RTX containing expression plasmid and tested for RTX expression by ELISA.

**Figure. 1.**
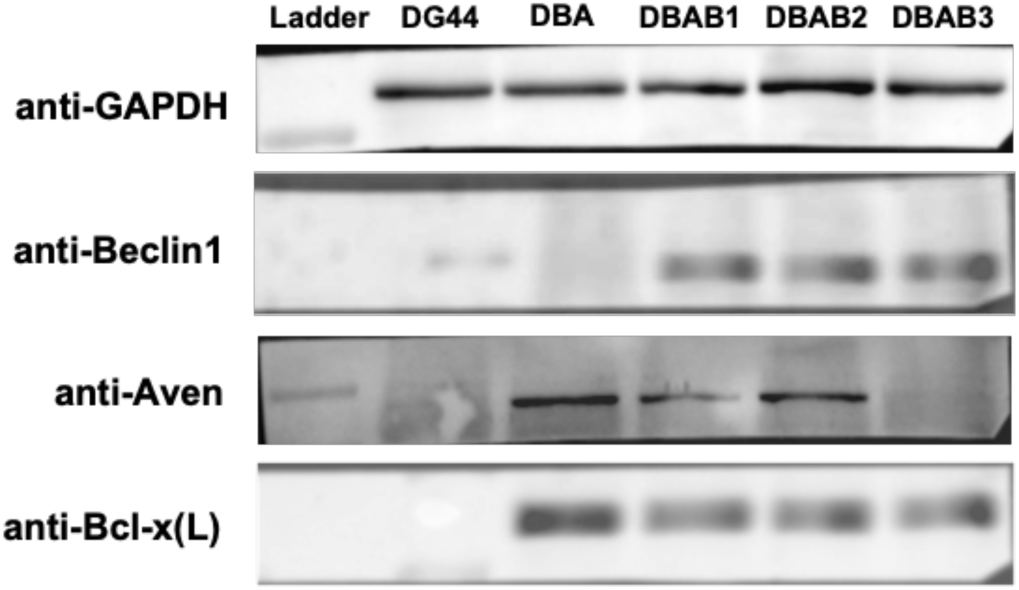
Western blots of lysates from control cells (DG44), cells transfected with p-BUD-Bcl-x(L)-Aven (DBA) and cells cotransfected with pBUD-BclxL-Aven and pcDNA3-BECN1 (DBAB 1-3) probed with antibodies anti-GAPDH and anti-Aven before stripping and with anti-Bcl-x(L) and anti-BECN1 post stripping.

### Quantitation of death fighting tendency of naïve and modified CHO cells in terms of apoptosis and autophagy

The death fighting tendency of modified cells was better during later hours of batch culture even when all essential media components got exhausted and there was an accumulation of byproducts like lactate and ammonia. The population of apoptotic cells in both control and modified cultures was measured using Annexin-V-PI based flow cytometry during early and late hours of batch culture. The population of apoptotic cells which was around 39% after 48 hours in the control culture increased to 72.5% in 96 hours **(Figure 2A and 2B)**. However, the modified cells performed better in terms of their tendency to fight apoptosis during both early and late hours of the culture, which can be seen clearly in Figure 2A where a negligible change in the apoptotic cell population was observed in 48 hours and 96 hours (25.0% and 23.6%). Similar trends were observed for percentage of apoptotic cells by dual acridine orange/ethidium bromide (AO/EB) staining **(Figure 2C)**. The autophagy assay showed a higher level of autophagy in control cells compared to modified cells. The higher band intensity of LC3-II at 96 hours compared to 48 hours in control cells indicates increased autophagosome formation in control cells during the later hours of batch culture **(Figure 2D)**. However, this activity was found to be balanced in modified cells, and there was a negligible change in LC3-II band intensities between 48 hours and 96 hours of culture. These results indicate that these host cell modifications were able to reverse the cell death due to apoptosis as well as autophagy.

**Figure 2.**
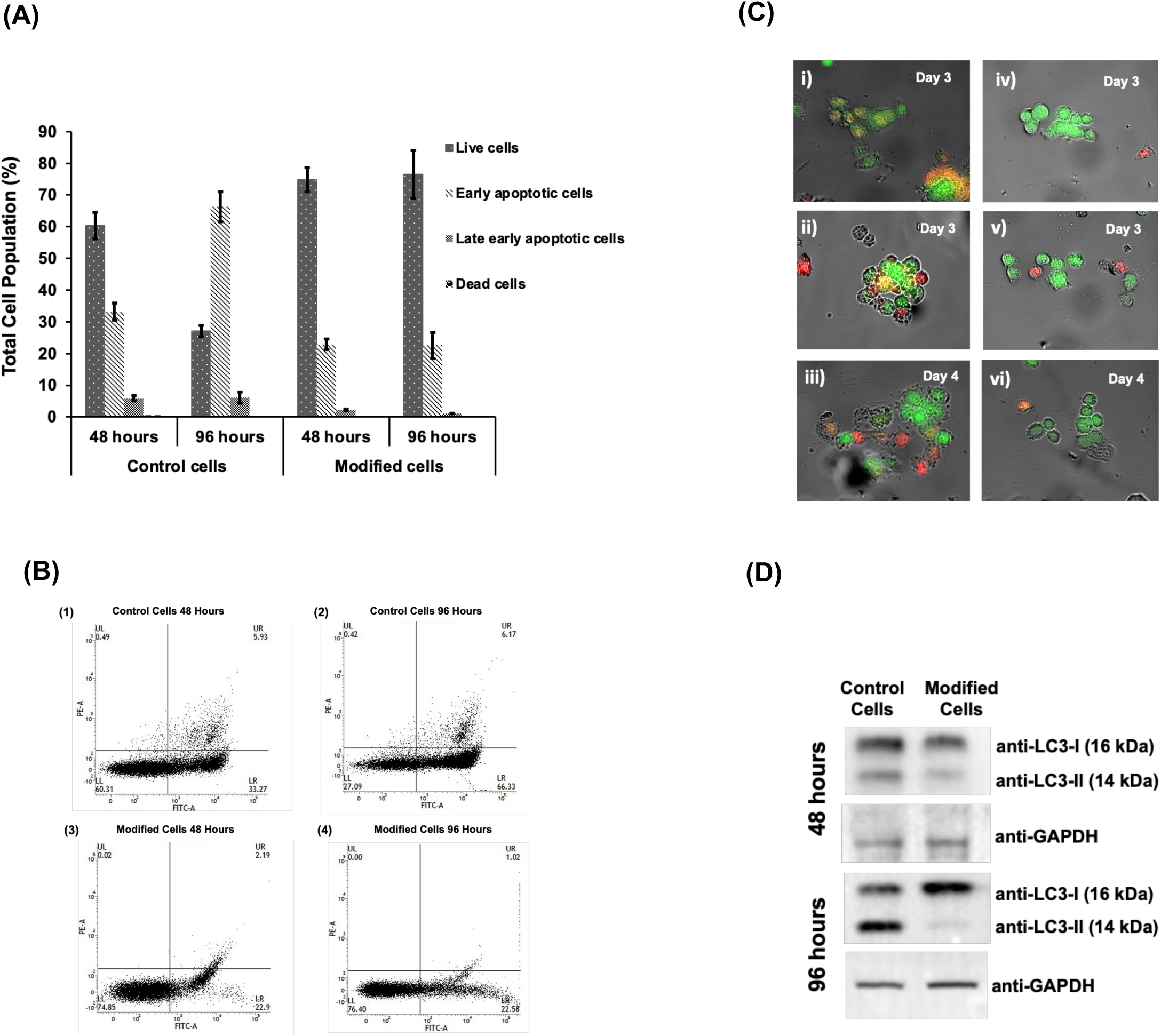
**(A)** Apoptosis quantification of control and modified cells using PI-Annexin V FITC based FACS assay (n= 3, mean ± SD). **(B)** FACS results of Annexin V-FITC and PI assay. Scatter diagrams of control cells during 48 and 96 hours of culture (1-2) and modified cells during 48 and 96 hours of batch culture (3-4). **(C) (i)** Normal DG44 cells on day 3 (early apoptotic cells). Nucleus showed yellow-green fluorescence by acridine orange (AO) staining and concentrated into a crescent or granular that located in 1 side of cells. **(ii)** Normal DG44 cells expressing RTX on day 3 and **(iii)** day 4. Late apoptotic and necrotic cells showing uneven orange-red fluorescence and an unapparent outline found individually as well as in lumps which is becoming dissolved or near disintegration on day 4. **(iv)** Modified cells on day 3. **(v)** Modified cells expressing RTX on day 3 and **(vi)** on day 4. The apoptosis delay found in modified cells as compared to the normal DG44 cells expressing RTX is in line with results of quantitative apoptosis analysis done by FACS. **(D)** Western blot showing the levels of LC3-I and LC3-II during 48 and 96 hours of culture. GAPDH was used as a loading control.

### Comparative growth and expression profile of naïve and modified CHO cells

The control and the modified cells with and without RTX genes were grown in T-25 flasks and their growth and expression profiles were monitored. It was observed that the untransfected control cells grew faster and better than the control cells transfected with RTX which showed a long lag phase. The possible reason for the delay in growth could be the cellular stress triggered by the expression of the recombinant protein. This cellular stress can also elicit responses which later lead to the activation of cell death mechanisms. This would explain the decline in integrated viable cell count (IVCC) and cell viability observed upon RTX expression in the control culture **(Figure 3A and 3B)**. The IVCC of the modified cells was significantly better than control. More importantly the IVCC did not decline upon RTX expression as had happened in the RTX expressing control cells. This could be ascribed to the over-expression of the BECN1 gene in the modified cells, which is known for promoting cell survival during nutrient deprived conditions (autophagosome formation); and is possibly involved in replenishing its nutrient supply. An integrated viable cell count of 4.1*10^6^ cells ml^-1^ was obtained for the modified system as compared to the DG44 control cells with an IVCC of 3.1*10^6^ cells ml^-1^. The modified cell line was able to sustain protein expression for seven days leading to a higher level of product formation. RTX expression thus increased by 7 - 7.5 folds in the modified cell line compared to the unmodified control **(Figure 3C)**.

**Figure 3.**
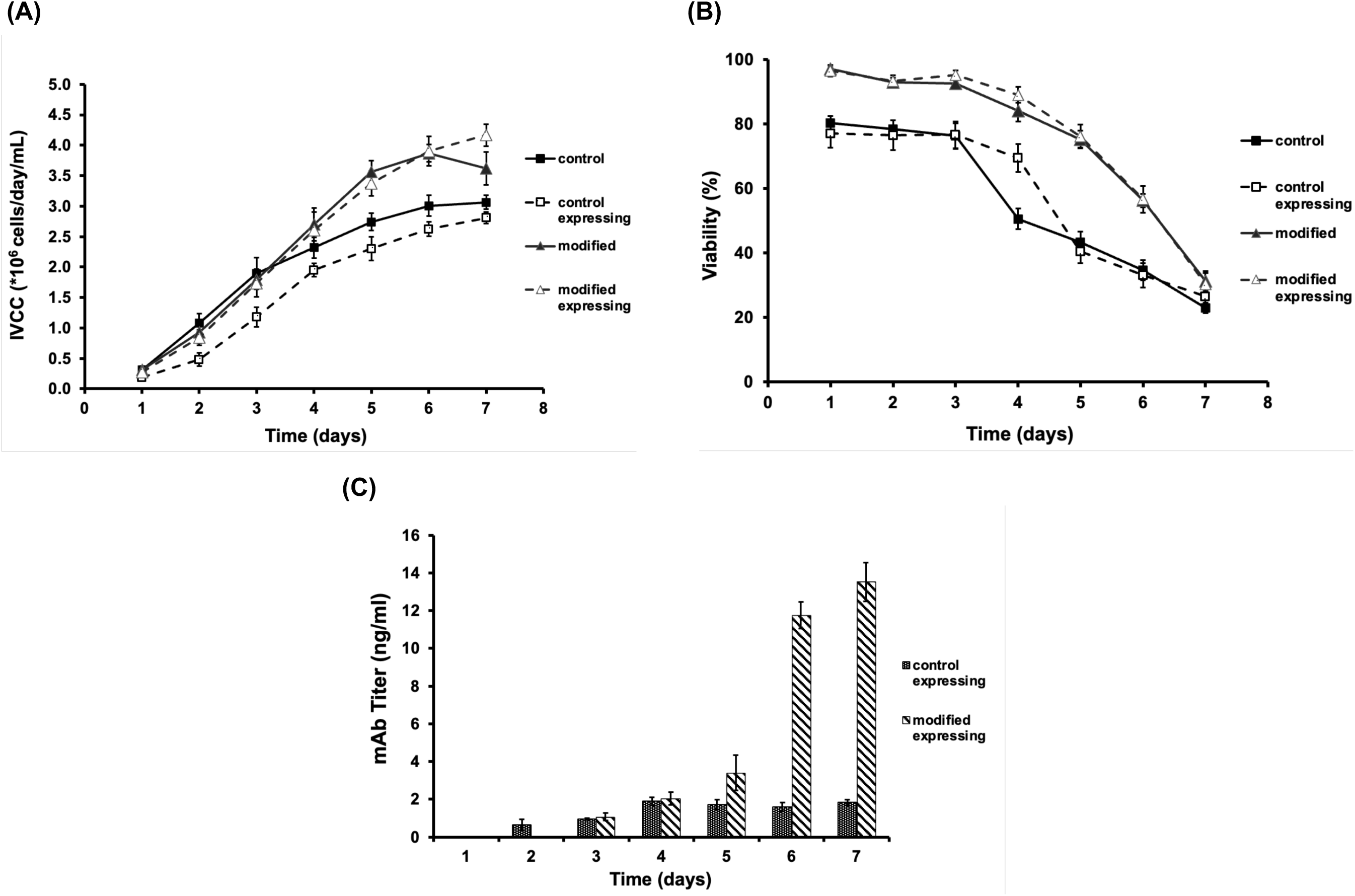
Modified cells are able to counter the stress associated with RTX expression. **(A)** Integrated Viable Cell Count (IVCC), **(B)** cell viability (%) and **(C)** production profile of control and modified cells expressing Rituximab (RTX) in static batch culture (n= 3, mean ± SD).

Both control and RTX expressing control cells showed nearly complete consumption of glucose by the third day. The residual glucose levels remained constant thereafter at a low value of 1 mM. Simultaneously there was accumulation of lactate up to 20 mM with the RTX expressing cells producing slightly elevated levels of lactate **(Figure 4)**. This can possibly be correlated with lower IVCC, since higher lactate accumulation in the RTX expressing cell line could have occurred due to cell death. The modified cells as well as its RTX expressing counterpart showed comparatively slower but more efficient utilization of glucose with residual glucose concentration falling to zero **(Figure 4)**. Lactate accumulation continued till the end of the glucose consumption phase in both cultures. However, unlike control the level of lactate with and without RTX expression was similar. These results clearly show that the modified cells were able to counter the stress associated with RTX expression.

**Figure 4.**
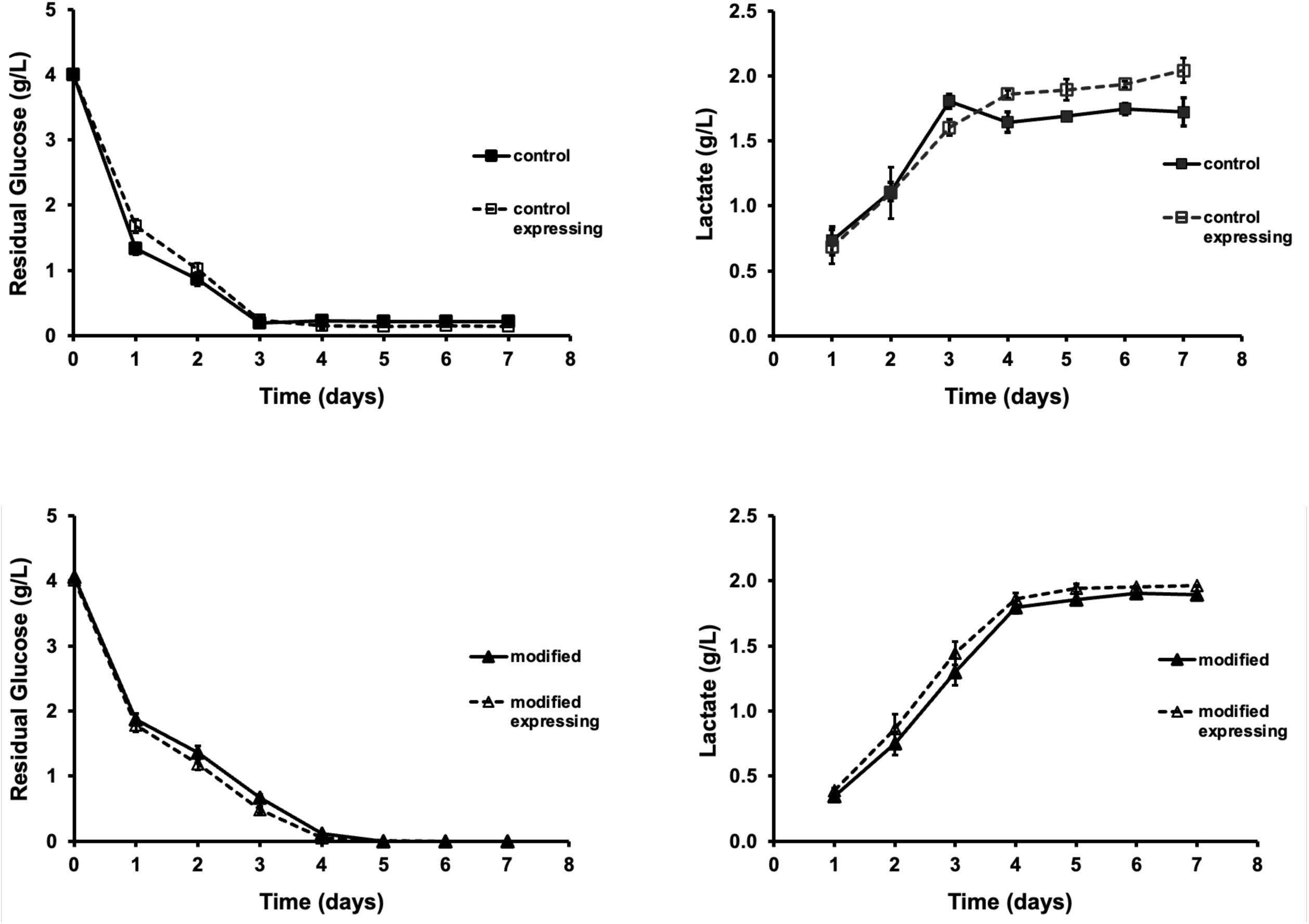
Lactate and glucose profiles of control and modified cells expressing Rituximab in static batch culture (n= 3, mean ± SD).

### Analysis of amino acid uptake

In order to identify the causes behind improved viability and expression capability of the modified cells, we decided to do amino acid profiling of the CHO cells grown in minimum essential media. Clearly substrate including amino acids uptake and/or secretion would have a direct bearing on cellular health. We were thus interested in identifying the critical nutrients whose differential uptake from the medium led to better performance. Since we were growing cells in minimal media where the concentration of the amino acids was fairly low, we observed a significant level of noise in the data while calculating uptake/secretion rates. We therefore used a least square fit over multiple time-points to estimate the rates of uptake or secretion of various amino acids from the medium. Since uptake rate would vary with culture conditions, we split the analysis into two broad phases based on nutrient uptake; phase I (from day 0 to day 3) where most of the glucose was taken up by the cells and phase II (from day 4 to day 7) where glucose consumption was negligible. This method gave us a reasonably good estimate of average uptake rates of different nutrients. Since the experiment was in a 2×2 format, we first estimated the uptake differentials between RTX expressing and non-expressing, control cells and then did a similar analysis on the modified cells (RTX expressing and non expressing cultures) **(Table 1A and 1B).** We observed that the amino acids levels in CHO DG44 control cells kept varying in the extracellular medium demonstrating that they were both secreted as well as taken up by the cells. This has been reported earlier^31–33^ and it is now well accepted that amino acid transport is bidirectional.

**Table 1A.**
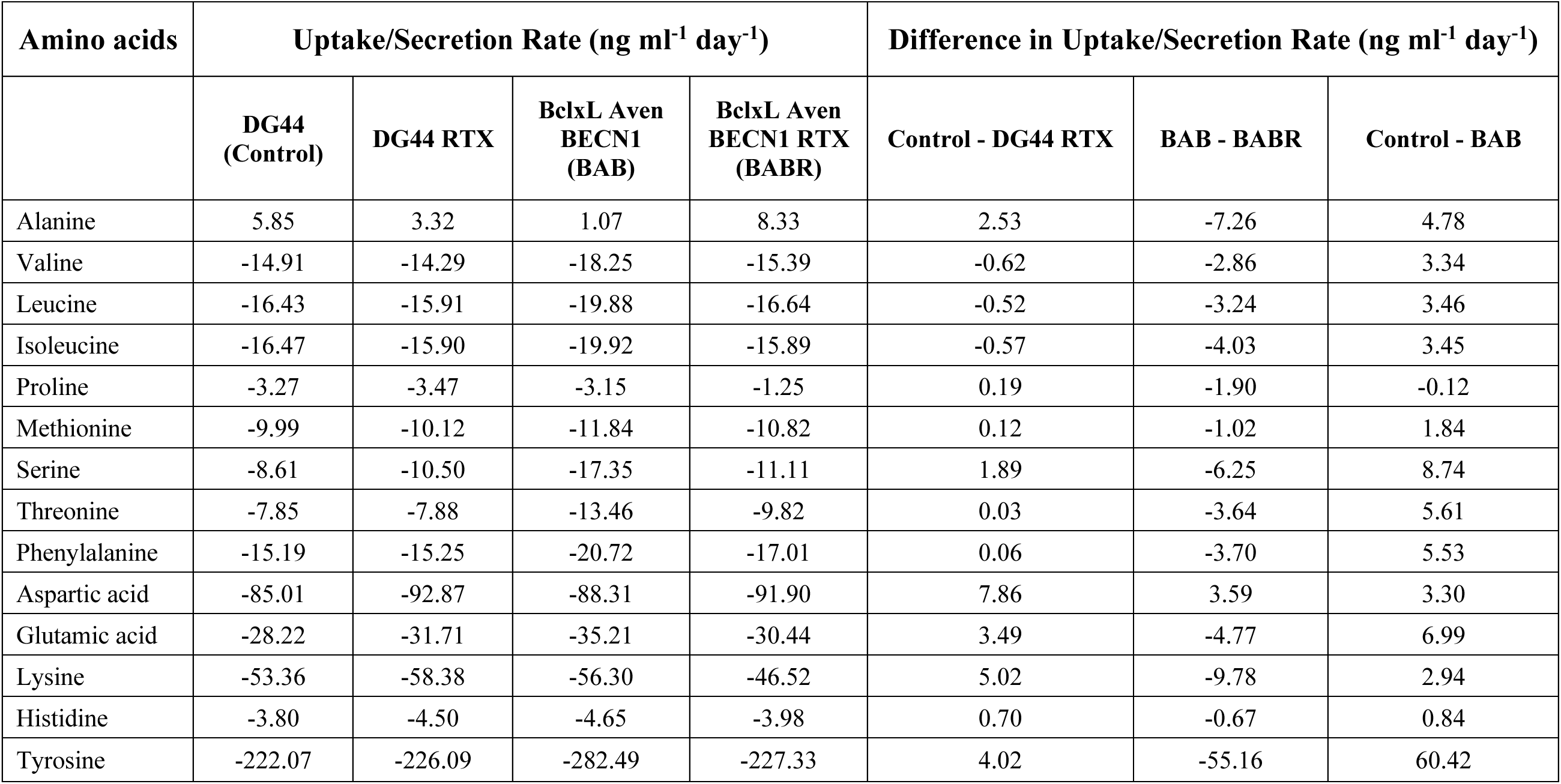
Differentials of amino acids uptake/secretion (From D0 to D3) for RTX expressing and non-expressing control and modified cells along with differences in amino acid uptake rates among different cell lines.

**Table 1B.** Differentials of amino acids uptake/secretion (From D4 to D7) for RTX expressing and non-expressing control and modified cells along with differences in amino acid uptake rates among different cell lines.

| Amino acids | Uptake/Secretion Rate (ng ml<sup>-1</sup> day<sup>-1</sup>) |  |  |  | Difference in Uptake/Secretion Rate (ng ml<sup>-1</sup> day<sup>-1</sup>) |  |  |
| --- | --- | --- | --- | --- | --- | --- | --- |
|  | DG44<br>(Control) | DG44 RTX | BclxL Aven<br>BECN1<br>(BAB) | BclxL Aven<br>BECN1 RTX<br>(BABR) | Control - DG44 RTX | BAB - BABR | Control - BAB |
| Alanine | -0.76 | 1.42 | 23.65 | 9.25 | -2.18 | 14.40 | -24.41 |
| Valine | -0.28 | 0.85 | 3.87 | 3.85 | -1.13 | 0.02 | -4.15 |
| Leucine | -0.19 | 0.74 | 4.15 | 3.70 | -0.93 | 0.45 | -4.34 |
| Isoleucine | -0.33 | 0.04 | 5.02 | 4.45 | -0.38 | 0.56 | -5.35 |
| Proline | -0.16 | 0.27 | 0.75 | 1.37 | -0.43 | -0.62 | -0.91 |
| Methionine | -0.19 | 0.49 | 0.78 | 1.13 | -0.68 | -0.35 | -0.96 |
| Serine | -2.42 | -0.76 | 8.48 | 6.92 | -1.66 | 1.56 | -10.90 |
| Threonine | -2.80 | -0.90 | 6.40 | 3.79 | -1.90 | 2.61 | -9.20 |
| Phenylalanine | -0.60 | 0.98 | 3.44 | 3.07 | -1.59 | 0.37 | -4.04 |
| Aspartic acid | -11.00 | 2.16 | -2.63 | -4.12 | -13.16 | 1.48 | -8.36 |
| Glutamic acid | -8.10 | 0.24 | -0.51 | -1.59 | -8.34 | 1.08 | -7.59 |
| Lysine | 1.73 | 1.70 | 3.85 | 4.40 | 0.03 | -0.55 | -2.12 |
| Tyrosine | -17.17 | 16.26 | 12.64 | 1.68 | -33.43 | 10.96 | -29.81 |

The differences in amino acid consumption rates during the first phase for both control and modified cells with and without RTX expression are shown in Table 1A. Control cells were able to utilize almost all amino acids except alanine from the extracellular medium during the first three days of batch culture. Tyrosine was consumed at a very high rate by control cells. The consumption of aspartic acid, glutamic acid and lysine was also high. Glutamine uptake rate from the extracellular medium was highest among all amino acids for both control and RTX expressing control cells. It is well known that glutamine is required in greater amounts (generally 4 to 6 mM) in cell culture than any other amino acid due to its central role in cell growth and metabolism^34^. However, we used comparatively lower amounts of glutamine (∼3 mM) in culture media order to minimize its accumulation and conversion into pyroglutamate and ammonia which are toxic for mammalian cell culture. RTX expression in control cells did not result in very significant changes in amino acid uptake rates during this phase of growth. The modified cells showed only a slight increase in consumption rates of all amino acids except for tyrosine whose uptake increased to 282 ng ml^-1^day^-1^ (27%) compared to control cells during this growth phase **(Table 1A)**. Glutamine levels were not detectable in the samples demonstrating that a complete consumption of glutamine took place in both RTX expressing and non-expressing modified cells within the first 3 days of batch culture. RTX expressing modified cells resulted in a negligible drop in uptake of all amino acids. However, the rate of tyrosine consumption dropped back to the level observed with control cells.

Table 1B shows the differences in amino acid consumption/secretion rates for control and modified cells again with and without RTX expression during phase II. RTX expression in control cells caused secretion of all the amino acids which were initially being consumed by these cells. However, the change from consumption to secretion or a major drop in consumption was most significant for tyrosine, aspartic acid and glutamic acid. This decline in the ability to take up amino acids (which is reflected in the negative values of the 5^th^ column in Table 1B) indicates that RTX expression in the control cells caused cellular stress and hence prevented them from utilizing these amino acids efficiently. This could be a possible reason behind the plateauing of recombinant protein levels in these cells. The amino acid analysis for the modified cells also showed a decline in consumption of most of the amino acids during this phase **(Table 1B)**. However, the modified cells expressing RTX continued to have a better uptake rate of most of the amino acids compared to non-expressing cells (which is reflected in the positive values of the 6^th^ column of Table 1B). A complete consumption of Histidine was observed in all cell lines except control cells during phase II. This clearly indicates that the cellular stress associated with RTX expression did not have a deleterious effect on the metabolic activity of the modified cells, which is reflected in the metabolome analysis by an increased uptake rate of amino acids. Amino acid uptake rates can be therefore considered as useful indicators of improved cellular health, which also affects the expression of recombinant proteins.

## Discussion

A problem that arises due to transient transfection in DG44 cells is that the product yields are lower compared to stable transfectants. Lower levels of expression could in turn lead to reduced cellular stress. We were therefore unsure whether our modified cell line, which was designed to counter the effects of stress, would indeed perform better than the control cell lines. However, from the data on cell viability, it is clear that RTX expression in control cells did lead to a significant level of cellular stress. If the level of RTX were higher, it is likely that the stress would be correspondingly higher. The performance of the modified cells under these circumstances would be expected to be equivalent if not better than what we obtained in the present study.

The main focus of the present work was to construct an improved host cell line so that it could be used as a model to study the phenotypic changes that correlate with improved expression levels. Given that the modified host had better cellular viability and could sustain RTX expression for a significantly longer time period made it a useful candidate for such a comparative analysis. Clearly the direct effect of the host cell modifications would be a down regulation of the apoptosis and upregulation of autophagy pathways. Thus Bclx(L) over expression would lead to suppression of both apoptosis and autophagy^35,36^. Aven is also an anti-apoptotic protein that inhibits caspase activation by Apaf-1 self-association. Aven binds to Bcl2, Bcl-x(L) and Mcl-1 enhancing the anti-apoptotic property of Bcl-x(L) following caspase-1 induced apoptosis^37^. To use the cell survival property of autophagy, BECN1 was introduced as it expresses the Beclin1 protein which binds with a class III phosphoinositide 3-kinase and initiates formation of the autophagosome thus enhancing autophagy^38^. All these effects are fairly well known and represent the specific and immediate effect of these modifications.

However, we were interested in locating the system level effects of these modifications and how these synergistically impact on cellular health. We hypothesized that a cascade of events would be triggered which are difficult to predict given the complex dynamics of cellular interactions. The final effect however would manifest itself in actual changes in critical metabolic pathways which improve cell viability and recombinant protein expression. We therefore studied the exometabolome which has no direct connection with the modifications introduced for blocking apoptosis and balancing autophagy pathways. It was very interesting to observe that RTX expression in control cells led to reduced amino acid uptake in the later phase of growth even though RTX build up had stopped. However, in the modified system RTX expression did not significantly change uptake rates. Rather the uptake of amino acid was slightly higher in the expressing cells compared to non expressing cells especially in the later stages of growth. Concomitantly RTX levels continued to rise, and these twin abilities of sustained expression and higher amino acids are clearly related. A stoichiometric analysis between these two will need a more detailed analysis of the kinetics of uptake and product formation. However, it is clear that this ability of the modified cells to handle the stress associated with RTX expression manifests itself in enhanced amino acid uptake and simultaneously sustained expression of the recombinant protein.

## Acknowledgements

We would like to thank Shirish Tripathi and Neetu Tyagi for extending their support in cell line maintenance. The support and funding received from Department of Science and Technology (DST) is gratefully acknowledged.

