## Supplemental Figures for "Correlating amino acid profiles with improved expression capability of superior CHO based expression platforms"

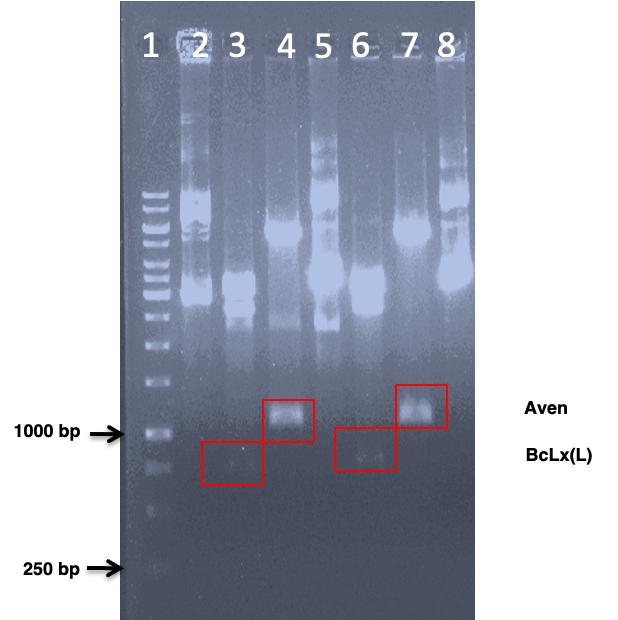

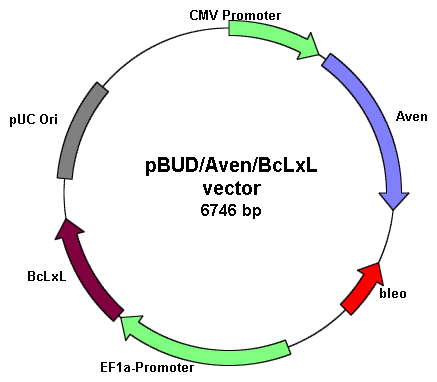

**(B)**

**(A)**

**(C)**

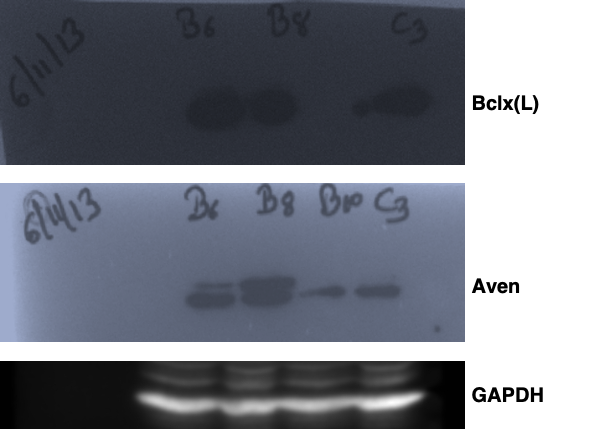

**Figure S1. (A) Final confirmation of pBUD Bcl-x(L) Aven** (Lane 1: Ladder; Lane 2: Construct 1; Lane 3: Bgl II-Not I digested construct 1; Lane 4: Sal I-Xba I digested construct 1; Lane 5: construct 2; Lane 6: Bgl II-Not I digested construct 2; Lane 7: Sal I-Xba I digested construct 2) **(B) pBUD- Bcl-x(L)-Aven construct (C) Western blot confirming expression of Bclx(L) and Aven in CHO cells** (GAPDH is used as a loading control).

Antibodies used: (1) Bclx(L) antibody: Bclx(L) mouse monoclonal IgG, Santa Cruz (concentration used 1:1000), cat. # sc-8392 (2) Aven antibody, cat. # 2300S (concentration used 1:1000) (3) Anti-GAPDH: Santa Cruz (concentration used 1:1000) (3) Secondary anti-rabbit IgG HRP linked antibody (concentration used 1:10000), cat. # 7074S) (4) Anti-mouse antibody: Goat anti-mouse IgG HRP (concentration used 1:10000), cat. # sc-2005.

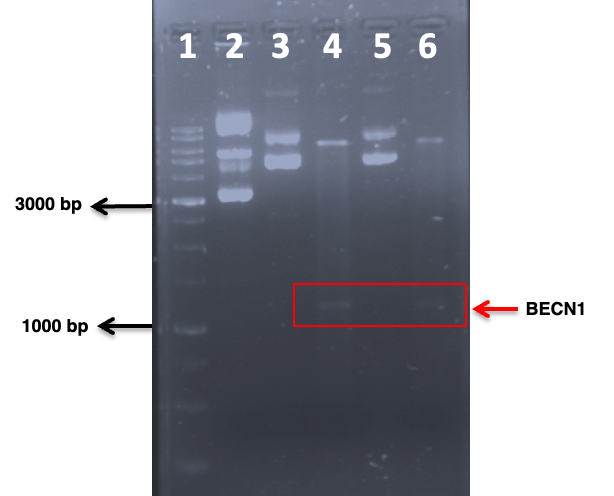

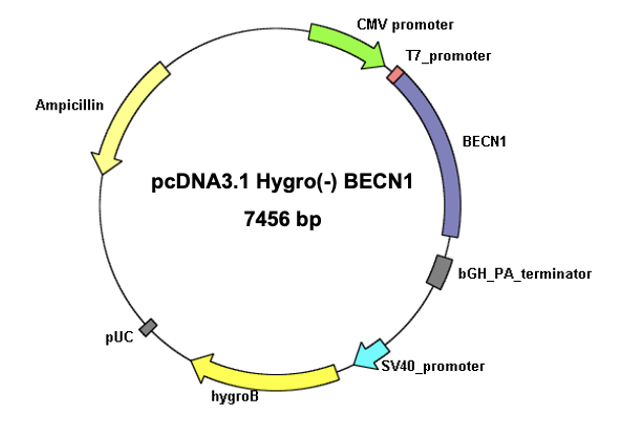

**(B)**

**(A)**

**(C)**

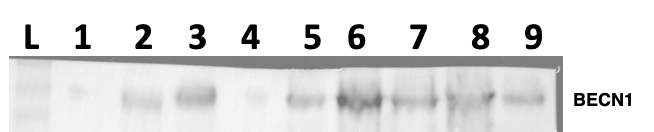

**Figure S2. (A) Ligation confirmation of BECN1 to pcDNA 3.1(-) Hygro by restriction digestion with Bam HI –Xba I** (Lane 1: Ladder; Lane 2: Plasmid backbone; Lane 3: construct 1; Lane 4: double digested construct 1; Lane 5: construct 2; Lane 6: double digested construct 2) **(B) pcDNA 3.1(-) Hygro - BECN1 construct (C) Westen blot confirming the expression of pcDNA3-BECN1 in CHO cells** (Lane 1: Ladder; Lane 2: DG44 cell lysate; Lane 3-9: lysates of transfectants).
Antibodies used: (1) Anti BECN1: BECN1 (H-300), Santa Cruz (concentration used 1:1000), cat. # sc-11427 (2) Anti GAPDH: Santa Cruz (concentration used 1:1000) (3) Secondary Anti rabbit IgG, HRP linked antibody, (concentration used 1:10000), cat # 7074S) (4) Anti Mouse antibody: Goat anti-mouse IgG HRP (concentration used 1:10000), cat. # sc-2005.

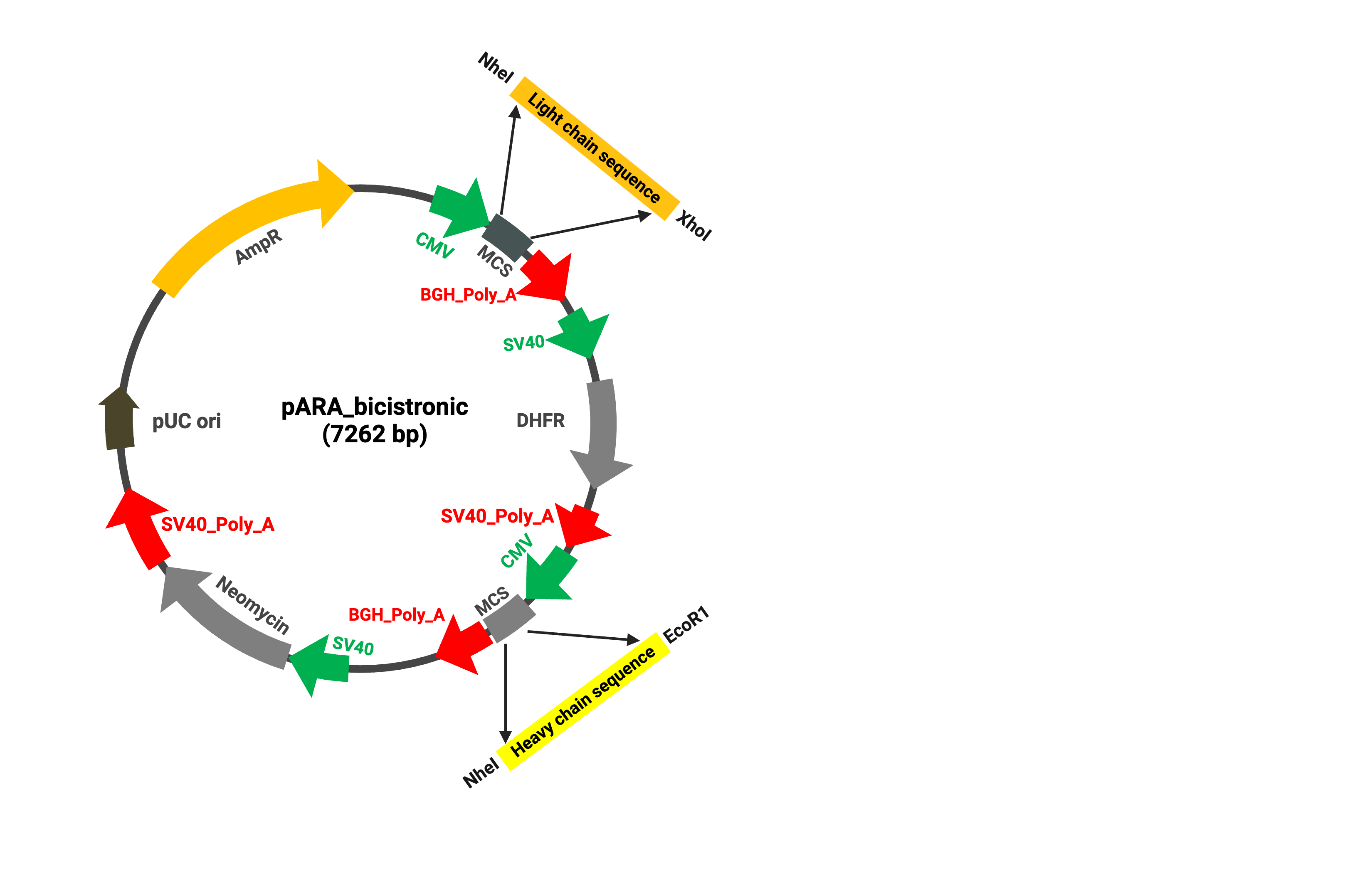

**Figure S3. Plasmid construct for Rituximab expression.**
